# What does domain-specific pretraining buy? A 170,000-parameter plant peptide language model against the ESM-2 scaling curve

**DOI:** 10.64898/2026.10.19.760098

**Authors:** KL Vomo-Donfack, M Aberbache, A Cantez, A Hoszu, G Ginot, VG Doblas, I Morilla

## Abstract

Small signalling peptides (SSPs) coordinate development and stress response in plants, and the computational tools that identify them represent sequences using general-purpose protein language models trained across all known proteins. How much a representation learned on the SSP sequence space itself is worth, measured against generalist embeddings of varying size, has not to our knowledge been quantified. We pretrain PhytoPepBERT, a 169,567-parameter masked language model implemented from scratch in NumPy with no deep-learning framework, on 1,469 plant SSP precursors retrieved from UniProtKB spanning five peptide families (RALF, PSK, CLE, PEP, RGF) and a length-matched negative class of non-SSP plant proteins. Sequences are clustered at 70% alignment identity and every cross-validation fold is grouped by cluster, so no homologue of a held-out sequence appears in training.

Domain pretraining is worth a great deal relative to where it starts. Family classification by *k*-nearest neighbours over frozen embeddings reaches 85.2% ± 2.4% against 58.0% ± 4.3% for an architecturally identical, identically initialised untrained backbone (chance 16.7%), and against 75.1% for dipeptide composition and 67.6% for BLOSUM62. The advantage is largest where the backbone stays frozen: linear probing gives 82.9% against 67.6% untrained, while full fine-tuning narrows the gap to 85.4% against 83.6%. Masked-residue accuracy is 20.5% against 3.4% untrained, elevated across all five peptide families and lowest on the non-SSP negatives.

Benchmarked against ESM-2 at four scales under an identical protocol, those 169,567 domain-pretrained parameters buy approximately what 7.5M generalist parameters buy: PhytoPepBERT ties `esm2_t6_8M_UR50D` (86.1% ± 2.8%, two of five paired folds) and is overtaken thereafter, by 2.5 points at 35M, 7.0 at 150M and 11.6 at 650M, losing every paired fold above the smallest checkpoint. TAPE, the second representation the reference pipeline uses, leads by 3.6 points at 92M. Generalist scale therefore continues to pay on this task well beyond the point where a domain model of this size can follow. We report an exchange rate rather than a victory: domain pretraining delivers roughly a forty-fold parameter efficiency against the generalist curve, and nothing beyond it.

## Introduction

Plant small signalling peptides (SSPs) are a large and functionally diverse class of intercellular messengers that regulate nearly every stage of development and much of the response to biotic and abiotic stress [1, 2, 3]. The families are defined as much by function as by sequence. CLAVATA3/ESR-related (CLE) peptides control stem-cell homeostasis in the shoot and root apical meristems and, as the twelve-residue TDIF peptide, suppress differentiation in the vascular cambium [5, 6, 7]. Rapid Alkalinization Factors (RALFs) arrest root growth and, through the FERONIA receptor kinase, couple cell-wall integrity to growth control [8, 9]. Phytosulfokines (PSKs) are sulfated pentapeptides that promote cellular proliferation [10]. Plant Elicitor Peptides (PEPs) are released from PROPEP precursors on wounding and activate innate immune signalling [11, 12]. Root Growth Factors, also called GOLVEN or CLEL peptides, maintain the root stem-cell niche and modulate auxin transport during gravitropism [13, 14]. Collectively these peptides constitute a signalling layer comparable in breadth to the classical phytohormones, and one whose membership is still being enumerated [4].

Enumerating it computationally is hard, for reasons that are intrinsic to the molecules. SSPs are translated as precursors, trafficked through the secretory pathway, proteolytically processed, and frequently modified post-translationally by tyrosine sulfation, proline hydroxylation or hydroxyproline arabinosylation before they are recognised by receptor kinases at the cell surface [1, 3]. The mature product is often shorter than fifteen residues. The precursor that carries it is short, frequently disordered outside the functional motif, and under weak selective constraint elsewhere. Primary-sequence conservation between orthologues is correspondingly poor, so homology search — the default instrument of comparative genomics — recovers only the most conserved families reliably, and a short conserved motif embedded in a variable precursor is precisely the configuration that profile methods handle least well. Dedicated resources [15, 16] and machine-learning classifiers [17] have become the practical route to genome-wide SSP discovery.

Those classifiers have converged on an architectural pattern inherited from protein machine learning at large. Since transformers [19] were adapted from natural language to biological sequence [20, 21], a succession of general-purpose protein language models — ESM-1b and ESM-2 [21, 22], ProtTrans [23], ProteinBERT [24], and generative counterparts such as ProGen and ProtGPT2 [25, 26] — have been trained by masked or autoregressive modelling over tens of millions of sequences spanning the tree of life. Their representations transfer remarkably well, and the standard recipe for a new task is now to embed with one of them and learn only a task-specific head. S^2^-PepAnalyst [17] follows exactly this pattern for plant SSPs, and does so twice over: it embeds each sequence independently with two protein language models — ESM-2 and TAPE [18] — refines both through a persistent-homology feature extractor, selects between them by reinforcement learning, and classifies the result with a convolutional network. Both representations are borrowed; only the classifier is learned on plant data.

This is a reasonable default, and for most protein tasks it is the right one. But it embeds an assumption that has not, for this domain, been tested. Plant SSPs occupy a narrow and atypical region of sequence space: short, compositionally skewed, often disordered, and dominated by a handful of processing and receptor-binding motifs embedded in otherwise unconstrained precursors. A model trained to represent the whole protein universe allocates very little of its capacity to that region. A representation learned on it specifically might therefore capture structure the generalist averages away. Equally, it might not: a few thousand sequences is a thin diet for a transformer, and the generic model may already encode everything the domain contains.

Evidence from adjacent fields does not settle the question; it shows that the answer is domain-dependent. In biomedical and scientific natural language processing, pretraining on in-domain text rather than fine-tuning a general model yields consistent and sometimes large gains [27, 28, 29], and in genomics a compact domain model has recently been argued to compete with much larger generalist alternatives [31]. Within peptide modelling, however, the dominant approach has been to fine-tune an existing general protein model rather than pretrain a new one — PeptideBERT [30], for instance, fine-tunes ProtBERT for hemolysis, solubility and non-fouling prediction. Whether domain pretraining pays in a domain as small and as specialised as plant SSPs is, to our knowledge, untested.

Answering it requires an evaluation one can trust, which turns out to be the harder half of the problem. Protein machine learning is unusually susceptible to two failure modes that inflate reported accuracy without any intent to deceive. The first is homology leakage: because biological sequences are related by descent, randomly partitioned folds place close relatives on both sides of the split, and the model is scored on near-copies of its training data. Avoiding it requires clustering by identity and splitting by cluster [32, 33, 34], and its neglect is a recognised contributor to irreproducibility in machine-learning-based science generally [35, 36]. The second is shortcut learning [37]: a model, or an evaluation, may exploit an incidental regularity — sequence length, amino-acid composition, or an invariant motif introduced by how a dataset was assembled — that correlates with the label but carries none of the intended signal. Both failure modes are easier to create than to notice, and a benchmark that admits either cannot support a claim about representation quality.

We therefore test the question as directly as we can, and construct the evaluation to resist both failure modes. We retrieve real plant SSP precursors from UniProtKB [40], pretrain a deliberately small transformer on them by masked language modelling, and compare the resulting frozen representation against ESM-2 and against non-learned baselines under one protocol, with the representation as the only variable. Five design choices follow from taking the comparison seriously rather than from convenience, and we state them up front because each of them lowers the numbers we report.

1. **Real sequences**. Every sequence is a retrieved UniProtKB entry. We previously ran this study on a corpus built by *in silico* mutagenesis from a handful of seed sequences, and report in Methods why that corpus could not support any conclusion about representation quality.
2. **Homology-aware evaluation**. Sequences are clustered at 70% alignment identity and cross-validation folds are grouped by cluster. On this corpus, ignoring clusters inflates accuracy by 3.8 to 8.6 points depending on the representation.
3. **A real negative class**. The non-SSP class is length-matched plant proteins retrieved from the same organisms and the same review status as the positives, not randomly generated strings.
4. **An architecturally identical control**. The principal comparison for the pretrained model is the same network with the same initialisation and the same seed, never trained. The difference between them is attributable to pretraining and to nothing else.
5. **A leakage control reported, not merely run**. A zero-parameter exact-substring classifier is reported alongside the learned methods.

We find three things. Domain pretraining works: PhytoPepBERT exceeds its own untrained initialisation by 27 accuracy points, and every non-learned baseline by at least ten. Measured against the generalist curve, those 169,567 parameters buy approximately what 7.5 M ESM-2 parameters buy — the two are level at the smallest checkpoint — but generalist scale keeps paying, and by 651 M the gap has opened to 11.6 points in ESM-2’s favour. What we report is therefore an exchange rate, not a victory. And the margin from pretraining is largest where a representation is used as a representation, with the backbone frozen, rather than where a downstream classifier can compensate for it.

## Results

### A corpus of real plant signalling peptide precursors

We retrieved 1,907 sequences from UniProtKB [40] (Table 1): five SSP families, identified by Pfam cross-reference where one exists (RALF, PF05498; PSK, PF06404) and by curated protein name otherwise (CLE, PEP, RGF), plus a negative class of plant proteins belonging to none of them. Sequences span 52–178 residues (median 105) across 278 organisms. The negative class was drawn to match the positives on organism and on Swiss-Prot/TrEMBL review status, so that neither taxonomy nor curation status correlates with the label.

**Table 1:** Corpus composition. All sequences retrieved from UniProtKB; none generated. Clustering at 70% alignment identity gives 1,163 clusters, and the train/validation split assigns whole clusters (1,469/438).

| class | *n* | identified by |
| --- | --- | --- |
| CLE | 400 | curated name, GOLVEN/RGF/CLEL excluded |
| PSK | 400 | Pfam PF06404 |
| RALF | 400 | Pfam PF05498 |
| NONSSP | 318 | organism- and status-matched non-SSP plant proteins |
| RGF | 301 | curated name (GOLVEN / CLEL / root meristem growth factor) |
| PEP | 88 | curated name (PROPEP / elicitor peptide) |
| total | 1,907 | 278 organisms, 52–178 residues, median 105 |

Before any model was trained we applied a leakage control, which we report rather than merely run. For each class we searched for any substring present in at least 90% of its members and in no member of any other class; such a substring would make the class label recoverable by exact matching and would invalidate any accuracy measured on the corpus. One family carries one: `DYIYTQ`, in 90.5% of PSK sequences. This is genuine biological conservation — the sulfated pentapeptide that defines the family — not an artefact of corpus construction, and we report it as a positive control: a representation that cannot separate PSK is failing at something easy. No other class carries a discriminator, and a zero-parameter classifier built from all of them scores 35.7% against 16.7% chance. The corpus is not solvable by exact matching.

### Pretraining learns residue-level structure

PhytoPepBERT is a three-layer, four-head pre-norm transformer with a model dimension of 64 and 169,567 parameters in total, implemented from scratch in NumPy with a hand-written autograd engine whose gradients agree with finite differences to a maximum absolute error below 3 × 10 ^4^ (Methods). It was pretrained by masked language modelling for 70 epochs on the 1,469 training sequences.

Validation masked-residue accuracy rose from 8.9% at epoch 1 to 23.3% at epoch 70 (Fig. 1a,b), and was still improving when we stopped: a linear fit over the final 25 epochs has a positive slope, and the gap between training (25.1%) and validation (23.3%) accuracy is small, so the model is undertrained rather than overfitted. The representation we go on to benchmark is therefore a conservative one.

**Figure 1:**
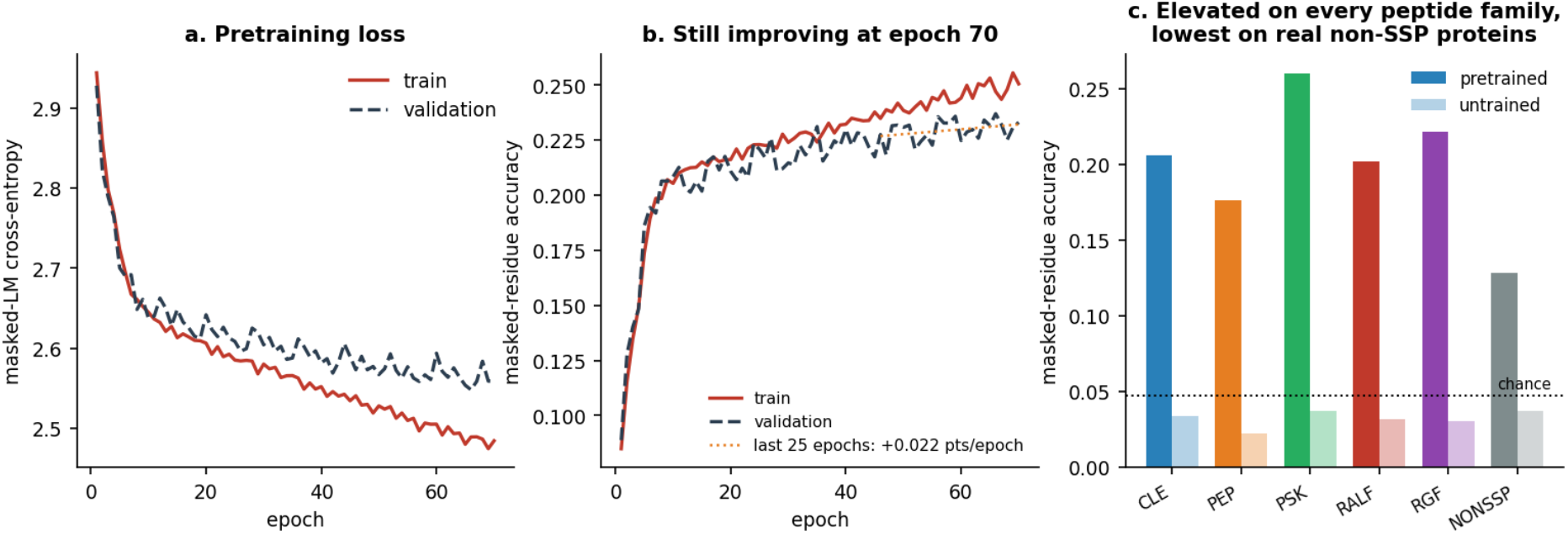
Pretraining. (a) Masked-language cross-entropy on training and held-out sequences over 70 epochs. (b) Masked-residue accuracy; the dotted line is a linear fit to the final 25 epochs, showing validation accuracy still rising when training was stopped. (c) Exact pseudo-perplexity by class for the pretrained and untrained backbones. All five peptide families are elevated; the non-SSP negatives are lowest. Chance is 1*/*21.

Scored by exact pseudo-perplexity — masking each position individually and predicting it from the rest of the sequence — the pretrained backbone recovers 20.5% of held-out residues against 3.4% for the untrained backbone, a factor of six (Fig. 1c). The margin is not uniform across classes, and the pattern is the one a reader should want: every peptide family is elevated (PSK 26.0%, RGF 22.1%, CLE 20.6%, RALF 20.2%, PEP 17.6%) while the non-SSP negatives sit lowest at 12.8%. The untrained backbone shows no such structure, scoring between 2.3% and 3.7% across all six classes. Pretraining has learned something specific to peptide precursors rather than general to plant proteins.

### The representation separates families far better than its untrained starting point

We evaluate the frozen representation by *k*-nearest-neighbour classification of family identity (*k* = 5) under five-fold cross-validation grouped by sequence cluster, so that no homologue of a test sequence appears in the training fold. The same protocol, the same folds and the same pooling are applied to every representation compared, so the representation is the only variable.

PhytoPepBERT embeddings classify the six classes at 85.2% *±* 2.4% (Fig. 2a, Table 2). The architecturally identical untrained backbone reaches 58.0% *±* 4.3%. That 27.1-point margin is attributable to pretraining alone: same parameter count, same depth and width, same initialisation, same seed, same evaluation.

**Table 2:** Family classification from frozen representations. Five-fold *k*NN (*k* = 5), folds grouped by 70%-identity cluster, mean *±* s.d. across folds. Six classes, chance 16.7%. The length-matched band spans 90–99 residues and holds three classes below fifteen members, so its column is reported for continuity rather than as support. Every learned representation above PhytoPepBERT is applied zero-shot.

| representation | params | full corpus ($n = 1907$) | length-matched ($n = 228$) |
| --- | --- | --- | --- |
| ESM-2 **t33_650M**, zero-shot | 651 M | **0.968 ± 0.012** | **0.939 ± 0.035** |
| ESM-2 **t30_150M**, zero-shot | 148 M | 0.921 ± 0.026 | 0.852 ± 0.075 |
| TAPE **bert-base**, zero-shot | 92 M | 0.887 ± 0.027 | 0.825 ± 0.041 |
| ESM-2 **t12_35M**, zero-shot | 33.5 M | 0.877 ± 0.030 | 0.800 ± 0.061 |
| ESM-2 **t6_8M**, zero-shot | 7.5 M | 0.861 ± 0.028 | 0.825 ± 0.072 |
| **PhytoPepBERT** | **169,567** | 0.852 ± 0.024 | 0.805 ± 0.054 |
| dipeptide composition | — | 0.751 ± 0.041 | 0.712 ± 0.095 |
| length + composition | — | 0.750 ± 0.049 | 0.708 ± 0.078 |
| BLOSUM62 | — | 0.676 ± 0.041 | 0.698 ± 0.065 |
| untrained backbone | 169,567 | 0.580 ± 0.043 | 0.673 ± 0.064 |
| length only | — | 0.310 ± 0.030 | 0.330 ± 0.069 |
| exact-substring classifier | 0 | 0.357 | — |

**Figure 2:**
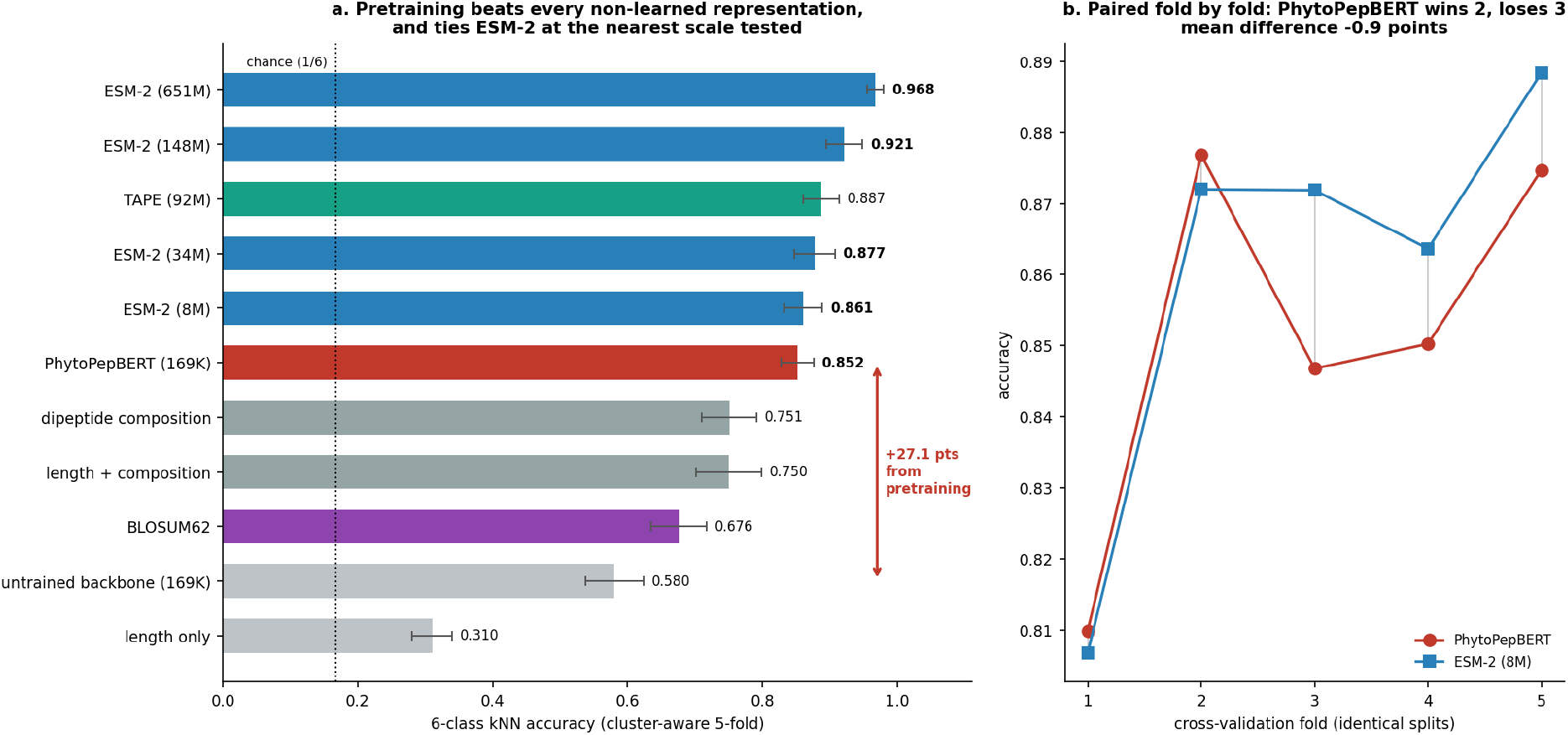
Representation comparison. (a) Six-class *k*NN accuracy from frozen representations under cluster-aware five-fold cross-validation. The bracket marks the margin between PhytoPepBERT and the architecturally identical untrained backbone, which is the quantity attributable to pretraining. Every learned comparator above PhytoPepBERT is applied zero-shot. (b) PhytoPepBERT against the nearest-scale ESM-2 checkpoint, scored fold by fold on identical splits: PhytoPepBERT leads in two folds and trails in three, the only comparison in the sweep that is not a clean loss.

It is also not reachable by cheaper means. Dipeptide composition, a 400-dimensional counting statistic, reaches 75.1%; length combined with single-residue composition 75.0%; mean-pooled BLOSUM62 substitution rows 67.6%; sequence length alone 31.0%. A domain-pretrained transformer with fewer than two hundred thousand parameters, trained on fewer than fifteen hundred sequences, encodes information about plant signalling peptides that substitution statistics and composition do not.

The same ordering holds within a length-matched band (90–99 residues, *n* = 228), where sequence length cannot act as a shortcut. We note that length was never a strong shortcut on this corpus — length alone classifies at 31.0% — and that three classes fall below fifteen members inside the band. We report the band for continuity with the comparable analysis in the nucleotide-domain literature and would not rest an argument on it.

### Against the ESM-2 scaling curve

The comparison this study was built to make is against the representation current SSP tools actually use. We embedded all 1,907 sequences with ESM-2 [22] at four scales — 7.5 M, 33.5 M, 148 M and 651 M parameters — and with TAPE [18], the second protein language model the reference pipeline runs in parallel with ESM-2. All are mean-pooled over final-layer per-residue representations exactly as PhytoPepBERT is, and evaluated on identical folds (Table 2, Fig. 3). The result is an exchange rate rather than a contest. At the smallest checkpoint the two are level: ESM-2 reaches 86.1% *±* 2.8% against PhytoPepBERT’s 85.2% *±* 2.4%, and read pairwise, PhytoPepBERT leads in two of five folds and trails in three (Fig. 2b). We report that as a tie, attaching no *P* -value: cross-validation folds share training data and are not independent observations [38, 39], and the fold-level record is reported descriptively throughout. Above the smallest checkpoint the curve rises monotonically and the tie does not survive: 87.7% at 35 M, 92.1% at 150 M and 96.8% at 650 M, a margin over PhytoPepBERT of 2.5, 7.0 and 11.6 points respectively, with every paired fold lost at every checkpoint above the smallest. TAPE, at 92 M parameters, reaches 88.7%, leading by 3.6 points and losing no fold.

**Figure 3:**
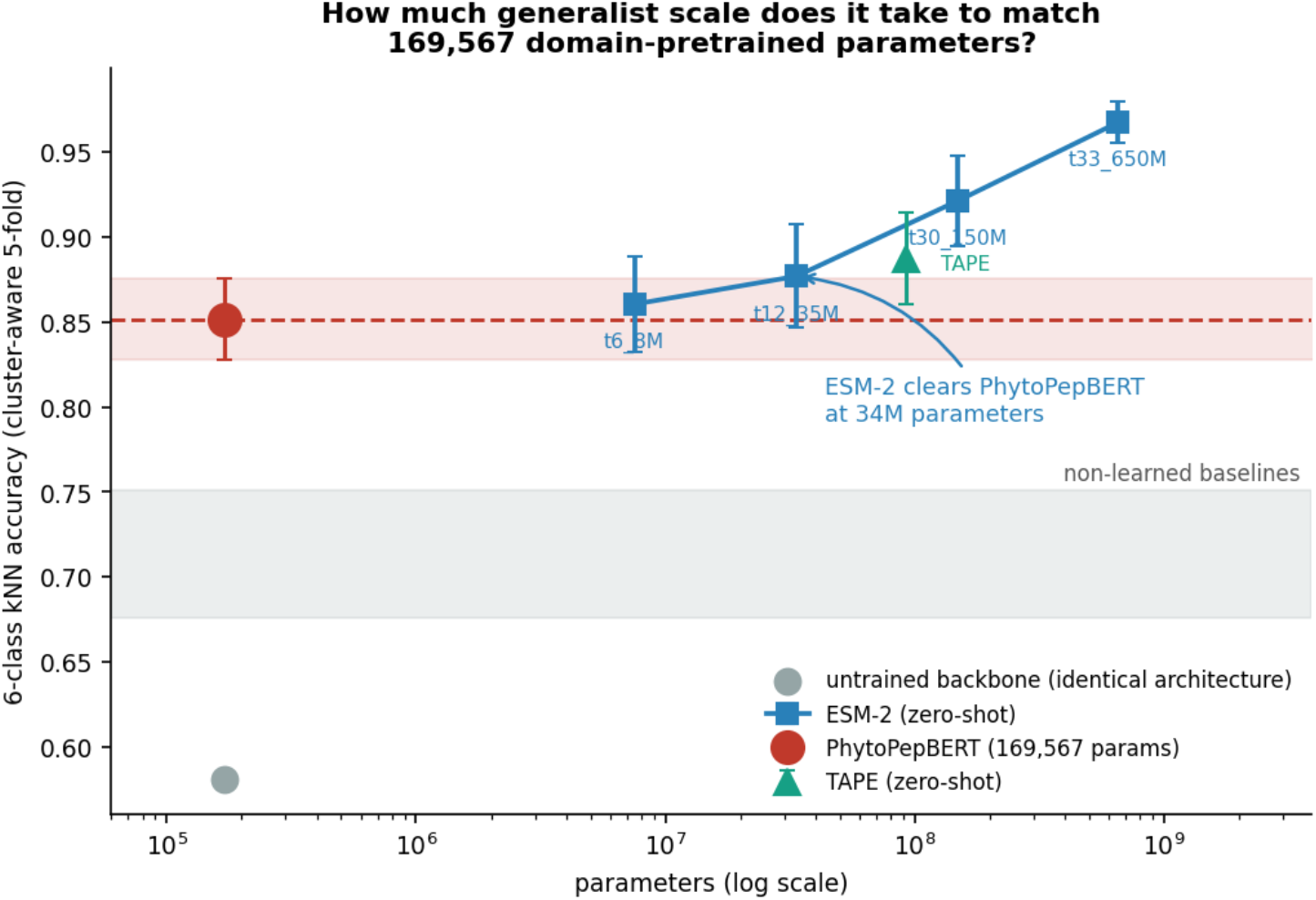
What domain pretraining is worth, measured against generalist scale. Six-class *k*NN accuracy against parameter count, all on identical cluster-aware folds. PhytoPepBERT (red, with fold s.d. band) sits level with the smallest ESM-2 checkpoint and is overtaken between 7.5 M and 33.5 M; the generalist curve continues to rise to 96.8% at 651 M. TAPE is shown at 92 M. The grey band spans the non-learned baselines, all of which both language-model families exceed. The untrained backbone, identical in architecture to PhytoPepBERT, is at bottom left.

Read as a scaling statement, 169,567 domain-pretrained parameters are worth approximately 7.5 M generalist ones on this task — a factor of about forty-four — and the curve crosses PhytoPepBERT somewhere between 7.5 M and 33.5 M. We do not have the resolution to locate that crossing more precisely, and would not read anything into the exact figure; what the data support is that the efficiency is large and that it runs out.

Two things bound the comparison, and we state them because they cut in opposite directions. The first favours us: ESM-2 and TAPE are applied zero-shot, having never encountered a plant SSP corpus, while PhytoPepBERT was pretrained on 1,469 of the 1,907 sequences the evaluation scores. Pretraining is self-supervised and uses no family labels, so the cross-validation remains honest with respect to the labels, but the representation is fitted to the evaluation corpus while its comparators are not. Measured directly, PhytoPepBERT scores 78.3% on the 438 sequences withheld from pretraining against 83.3% on matched samples of those it saw — a five-point transductive advantage. BLOSUM62, which saw nothing, differs by

1.5 points across the same subsets, so the effect is not an artefact of the held-out set being easier. Correcting for it would move the crossing point downward, not upward.

The second cuts against us, and matters for how the result should be read in context. The checkpoint that beats PhytoPepBERT most decisively, 651 M, is the one the reference pipeline actually uses: S^2^-PepAnalyst reports 1280-dimensional ESM-2 embeddings [17], which identifies that model. A domain representation of this size is therefore not a drop-in substitute for the embedding a production tool depends on, and we make no such claim.

### Pretraining matters most when the backbone is frozen

Frozen embeddings are one way to interrogate a representation; attaching a classifier is another, and the two answer different questions. We fine-tuned both backbones under two regimes: a linear probe over frozen weights, and full end-to-end fine-tuning (Fig. 4).

**Figure 4:**
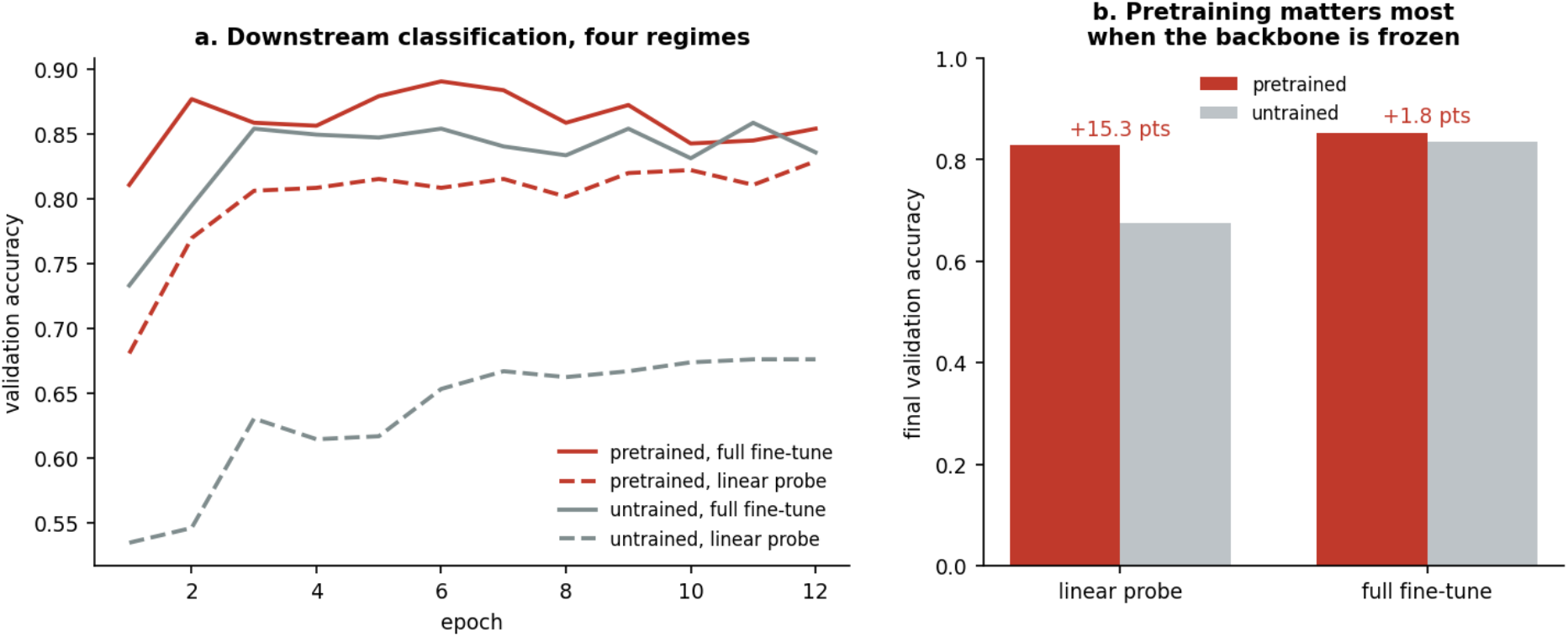
Downstream classification. (a) Validation accuracy over twelve epochs for both backbones under both regimes. (b) Final accuracy by regime. Pretraining is worth 15.3 points with the backbone frozen and 1.8 points end to end.

With the backbone frozen, pretraining is worth 15.3 points: 82.9% against 67.6% for the untrained network. With the whole network trainable, it is worth 1.8 points: 85.4% against 83.6%. The pretrained model also converges faster, reaching 89.0% at epoch 6 where the untrained network peaks at 85.8% at epoch 11.

We read this as a statement about corpus size rather than about pretraining. At 1,469 sequences a three-layer transformer can be trained to this task directly, and the pretrained weights buy a better frozen representation and faster convergence rather than a higher ceiling. Whether that gap widens on a corpus an order of magnitude larger is an empirical question this study cannot answer. It does, however, bear on the practical case: a tool that embeds sequences once and reuses the representation across tasks operates in the frozen regime, where the margin is fifteen points, not two.

### What the model attends to

Two analyses ask whether the learned structure is interpretable rather than merely useful (Fig. 5). Gradient saliency for masked-residue prediction, computed as the *L*_2_ norm of the gradient with respect to each input token embedding, concentrates on the masked positions: all seventeen masked positions in the example sequence score above the sequence-wide mean, and four are among the eight highest-scoring positions of 180. The model uses the positions whose prediction it is scored on — a sanity check rather than a finding, but a necessary one.

**Figure 5:**
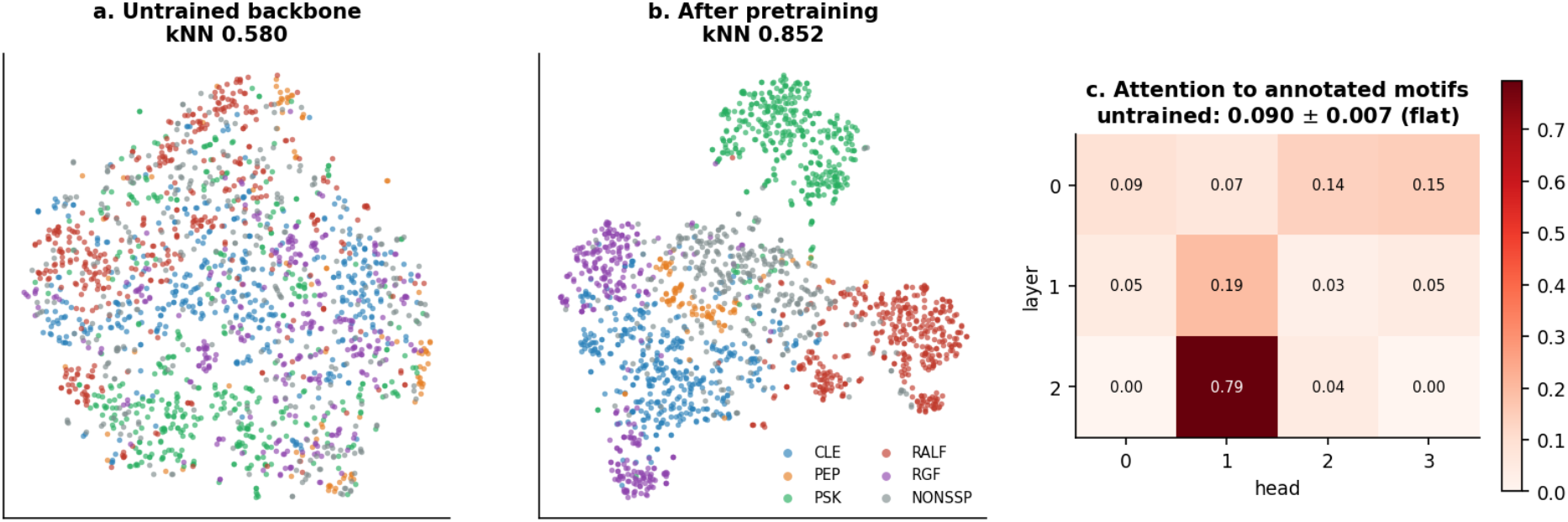
Embedding geometry and attention. (a,b) t-SNE [50] projections of pooled embeddings before and after pretraining, coloured by class, with the corresponding *k*NN accuracy. (c) Attention directed at curated mature-peptide annotations, by layer and head, after pretraining; the untrained backbone is flat at 0.090 *±* 0.007 across all twelve cells.

Attention directed at curated mature-peptide annotations is strongly head-specific. Averaged over query positions and sampled sequences, the untrained backbone distributes attention almost uniformly across all twelve layer–head pairs (mean 0.090, s.d. 0.007, range 0.083–0.107). After pretraining, the same twelve cells span nearly the entire available range (mean 0.133, s.d. 0.208), with one head in the final layer directing 79.4% of its attention to annotated motif positions while others fall to essentially zero. Pretraining induced specialisation that did not exist at initialisation.

This analysis carries a caveat we state plainly: only 2% of the SSP entries in the corpus carry curated `PEPTIDE` or `PROPEP` feature annotations in UniProtKB, because the corpus is drawn overwhelmingly from TrEMBL rather than Swiss-Prot. The attention result therefore rests on a small, better-annotated subset and should be read as suggestive.

## Discussion

The central result is the margin over the untrained backbone. PhytoPepBERT’s embeddings separate six classes at 85.2% where an architecturally identical network with identical initialisation, differing only in having never seen a peptide, reaches 58.0%. Those 27 points are attributable to pretraining and to nothing else, and they are not reachable by cheaper means: dipeptide composition reaches 75.1%, BLOSUM62 67.6%, length alone 31.0%. A 169,567-parameter masked language model trained on fewer than fifteen hundred real precursors learns something about plant signalling peptides that substitution statistics and composition do not encode.

Where that structure lives is visible in the fine-tuning results, and the pattern there is more informative than the headline number. With the backbone frozen, pretraining is worth 15.3 points; with the whole network trainable, 1.8. The representation matters most precisely when the downstream classifier cannot compensate for it, and end-to-end training on this corpus substantially rediscovers what pretraining supplied. This bears on how such a model would be used: a pipeline that embeds once and reuses the representation sits in the frozen regime.

The head-to-head does not favour us, and the shape of the result is more informative than the verdict. Against the smallest ESM-2 checkpoint, 7.5 M parameters, the two are level and we call it a tie. Against every larger one the tie disappears, monotonically and without a single fold won: 2.5 points behind at 35 M, 7.0 at 150 M, 11.6 at 651 M. TAPE, at 92 M, is 3.6 ahead. Read as a scaling statement rather than a contest, 169,567 domain-pretrained parameters purchase roughly what 7.5 M generalist ones do on this task, and the crossing lies between 7.5 M and 33.5 M. That is a parameter efficiency of around forty fold, and it is bounded: it does not extend, and generalist scale continues to deliver long after a domain model of this size has stopped.

We think the bounded version is the more useful finding. “A small domain model ties a generalist” would have been a claim about one comparison; an exchange rate is a quantity a reader can apply. It says that if the budget is a laptop CPU and no framework dependency, domain pretraining recovers the performance of a generalist two orders of magnitude larger — and that if the budget extends to a 651 M-parameter embedding, nothing we built competes with simply using it. Both halves are actionable, and the second is the one a practitioner choosing a representation today most needs to hear.

Two asymmetries qualify it further. PhytoPepBERT carries a measured five-point transductive advantage from having been pretrained on 77% of the evaluation sequences, so the true crossing point is lower than where we observe it. And the checkpoint that beats us by the largest margin is the one the reference pipeline uses, so the gap we report is the operationally relevant one rather than a worst case we constructed. Neither works in our favour.

What survives untouched is the comparison against our own starting point. The 27-point margin over an architecturally identical, identically initialised untrained network is unaffected by anything in the scaling analysis: it is measured at fixed parameter count, fixed seed and fixed evaluation, and it says that pretraining on fewer than fifteen hundred real precursors teaches a transformer something about plant signalling peptides. That finding does not depend on how it compares with a model trained on four thousand times more data.

The question we would put next is not whether a small domain model can replace a generalist embedding — on this evidence it cannot, above the smallest checkpoint — but whether the two are complementary, given that they are trained on different objectives at corpus scales separated by five orders of magnitude. Concatenating a 169,567-parameter domain representation with a 651 M-parameter generalist one costs almost nothing on top of the generalist already being computed, and the protocol above specifies the experiment completely.

## Limitations

### Corpus composition

Only 2% of the corpus is Swiss-Prot reviewed, so annotation quality is uneven and the attention analysis rests on a small subset. Although 278 organisms are represented, retrieval is weighted toward well-sequenced genera, and the taxonomic breadth of the result is narrower than the species count suggests. PEP is represented by 88 sequences against 400 for the largest classes.

### The scaling curve is four points

We resolve that the crossing lies between 7.5 M and 33.5 M parameters but not where within that interval, and a single corpus and task cannot establish that the exchange rate generalises to other peptide families or other domains. We also did not vary the domain model’s own size, so we cannot say whether a larger PhytoPepBERT would shift the crossing or merely overfit 1,469 sequences.

### Undertrained

Validation masked-residue accuracy was still rising at epoch 70, so the representation benchmarked here is a conservative estimate of what this architecture reaches on this corpus.

### Family assignment

Three of five families (CLE, PEP, RGF) were retrieved by curated protein name rather than Pfam domain, because Pfam does not model them as domains. We excluded GOLVEN/RGF/CLEL entries from the CLE query explicitly — UniProt names them “CLAVATA3/ESR (CLE)-related” for historical reasons although they constitute a separate family — and promoted them to their own class. Other name-based misassignments may remain.

### The length-matched band is weak

At *n* = 228 with three classes below fifteen members it carries little statistical weight. We report it for continuity, not as support.

## Conclusion

Domain-specific pretraining on a small corpus of real plant signalling peptide precursors produces sequence representations that exceed every non-learned alternative tested, and exceed an architecturally identical untrained network by 27 accuracy points. Measured against general-purpose protein language models at four scales, those 169,567 parameters are worth approximately 7.5 M generalist ones: level with the smallest ESM-2 checkpoint, overtaken by 33.5 M, and 11.6 points behind the 651 M model that the reference pipeline for this task actually uses. The contribution is therefore a quantity rather than a ranking — roughly a forty-fold parameter efficiency, and a clear limit to it. For a group whose constraint is compute rather than accuracy, a model that trains on a laptop CPU in NumPy and recovers the performance of a generalist two orders of magnitude larger is a usable tool. For a group whose constraint is accuracy, the generalist curve has not flattened by 651 M parameters and the right choice is to embed with it.

## Methods

### Corpus retrieval

Sequences were retrieved from the UniProtKB REST API [40], restricted to Viridiplantae (`taxonomy_id:33090`) and to 52–178 residues. RALF and PSK were identified by Pfam [41] cross-reference (`xref:pfam-PF05498` and `xref:pfam-PF06404`); CLE, PEP and RGF by curated protein and gene name, since Pfam does not model these families as domains. The CLE query explicitly excludes GOLVEN, root meristem growth factor and CLEL entries, retrieved separately as RGF. Retrieval was capped at three sequences per organism per family to limit taxonomic skew. The negative class comprises plant proteins matching none of the family queries, retrieved per organism to match the positives on both organism and Swiss-Prot/TrEMBL review status; organism-matched querying also avoids the whole-database scan a single broad exclusion query would require.

### Redundancy reduction and splitting

Sequences were clustered by single-linkage agglomeration at 70% global alignment identity (Needleman–Wunsch [43], BLOSUM62 [42], gap open 11, extend 1, via `parasail` [45]), with a 5-mer inverted-index prefilter, giving 1,163 clusters from 1,907 sequences. Whole clusters, not individual sequences, were assigned to the training (1,469) and validation (438) splits, stratified by family. We verified the outcome rather than trusting the procedure: across a sample of validation sequences the maximum identity to any training sequence is 0.70, and no validation sequence retains a relative above the clustering threshold. CD-HIT or MMseqs2 at the same threshold may be substituted.

### Leakage control

For each class we searched for any substring present in 90% of its members and in no member of any other class, scanning substrings of each class’s shortest member (any motif shared by a whole class must be a substring of its shortest sequence). A zero-parameter classifier built from the resulting motifs is reported alongside the learned methods. We adopted this control after discovering that an earlier, synthetically augmented version of this corpus — built by *in silico* mutagenesis from four seed sequences with each family’s conserved motif held invariant — was solved at 100% by exactly such a classifier, above every learned method we had benchmarked on it, including a protein language model. No accuracy measured on that corpus could support a claim about representation quality, and none of its results are reported here. We state this because the failure mode is easy to reproduce and hard to notice.

### Architecture

A pre-norm transformer encoder: 3 layers, 4 attention heads, model dimension 64, feed-forward dimension 256, GELU activations [52] in a pre-norm arrangement [53], learned positional embeddings, maximum sequence length 180, and a vocabulary of 26 (20 amino acids, `X`, five special tokens). Total parameters 169,567, including both the masked-language and classification heads. Implemented from scratch in NumPy with a hand-written reverse-mode autograd engine; every operation’s gradient was verified against central finite differences with a maximum absolute error below 3 × 10^−4^.

### Pretraining

Masked-language modelling with 15% of positions selected, of which 80% are replaced by `[MASK]`, 10% by a random amino acid and 10% left unchanged [20]. Adam [51], learning rate 4 × 10^−3^, batch size 16, 70 epochs, seed 0. The untrained control is the same architecture instantiated with the same seed and never trained; because the classification head is constructed last, the backbone and masked-language head are bit-identical between the two models.

### Pseudo-perplexity

Each non-special position of each validation sequence is masked individually and predicted from the remainder, up to 40 positions per sequence, reported as the fraction of masked residues recovered exactly.

### Embedding analysis

Pooled embeddings are the mean of final-layer hidden states over non-special, non-padding positions. Baselines: dipeptide frequency (400-dimensional, *L*_1_-normalised), sequence length alone, length with single-residue composition (21-dimensional), and mean-pooled BLOSUM62 rows (20-dimensional), the matrix loaded from Biopython [44] rather than transcribed by hand. All representations are pooled identically, so the comparison differs only in the representation.

### Cross-validation

Five-fold `StratifiedGroupKFold`, shuffled, seed 0, with each sequence’s cluster identifier as its group; *k*-nearest neighbours with *k* = 5; standardisation fitted on the training fold only. Grouping is not optional: scoring the same representations with ungrouped `StratifiedKFold`inflates accuracy by 3.8 points (dipeptide composition) to 8.6 points (BLO-SUM62). All results were produced with scikit-learn 1.3.2 [48], and we record the version because `StratifiedGroupKFold`’s fold-balancing behaviour differs between releases: on this corpus, 1.3.2 yields fold sizes of 400, 349, 359, 334 and 407, where 1.8.0 balances them to 382, 382, 381, 381 and 381. Fold compositions and therefore per-fold accuracies are version-dependent, and mean accuracies move by one to two points between the two. Conclusions do not, but exact reproduction requires the pinned version.

### Transductive-advantage measurement

PhytoPepBERT is pretrained on 1,469 of the 1,907 sequences the *k*NN evaluation scores, while ESM-2 and BLOSUM62 are zero-shot on all of them. To size this asymmetry we scored the 438 sequences withheld from pretraining separately and compared against matched random subsets of the 1,469 it saw, across five cross-validation seeds each. BLOSUM62, which saw nothing, serves as the control separating a genuine transductive effect from any intrinsic difference in subset difficulty.

### Fine-tuning

A linear classification head over the pooled `[CLS]` representation, 12 epochs, Adam. Full fine-tuning uses learning rate 1.5 *×* 10^−3^;the linear probe, with the backbone frozen, uses 8 *×* 10^−3^. Both are evaluated on the cluster-aware validation split.

### ESM-2 head-to-head

ESM-2 embeddings come from the `esm2_t6_8M_UR50D` checkpoint (6 layers, 320 dimensions, 7,512,474 parameters) loaded through `fair-esm` 2.0.0 under PyTorch 2.2.2, taking final-layer per-residue representations and mean-pooling over sequence positions only, excluding start- and end-of-sequence tokens, identical to the pooling used for PhytoPep-BERT:

~~~
_, _, toks = bc([(“q”, seq)])
=with torch.no_grad(): out = model(toks, repr_layers=[6])
emb = out[“representations”][6][0, 1:len(seq)+1].mean(0)
~~~

Embedding and analysis run as separate processes: PyTorch and NumPy/scikit-learn link different OpenMP runtimes, which on some platforms segfaults when both are loaded into one process.

### Explainability

Saliency is the *L*_2_ norm of the gradient of the masked-language loss with respect to each input token embedding. Attention-to-motif is the attention mass each (layer, head) pair directs at positions inside curated UniProtKB `PEPTIDE`or `PROPEP`feature ranges, averaged over query positions and over sampled sequences per family. Only entries carrying such annotations contribute.

### Software and reproducibility

All stochastic procedures use seed 0. PhytoPepBERT itself requires only NumPy, SciPy, scikit-learn and Matplotlib [46, 47, 48, 49]; corpus construction additionally requires `requests`, `biopython`and `parasail`, and the ESM-2 arm requires `torch` and `fair-esm`with network access to a model hub. Pretraining checkpoints every eight epochs and re-seeds its masking RNG at each call, so exact weight reproduction requires the same interruption pattern; trained checkpoints are provided so that no downstream result depends on reproducing them.

## Data and Code Availability

All code, the retrieval script, the exact corpus with cluster assignments and splits, trained weights, and the JSON outputs underlying every figure are available at https://github.com/MorillaLab/PhytoPepBERT. The corpus is reconstructible from the UniProtKB accessions recorded in the released `splits.json`.

## Acknowledgments

We gratefully acknowledge funding from the ‘Proyecto de Jóvenes Investigadores’ (UMA B1-2022-29 to I. Morilla), Consejería de Universidades, Ciencias y Desarrollo, fondos FEDER de la Junta de Andalucía (ProyExec_0499 to I. Morilla), MINISTERIO DE CIENCIA E INNOVACIÓN (PID2020-113378RA-I00 to V.G. Doblas).

